# Meiotic null *MSH2* and *SGS1* alleles in *S. cerevisiae* x *S. uvarum* hybrids result in near-haploid offspring with mixed parental chromosomal inheritance

**DOI:** 10.1101/2025.06.02.657486

**Authors:** Taylor K. Wang, Maitreya J. Dunham

**Author notes:** Correspondence to: Maitreya J Dunham.

## Abstract

In the natural world, genetic isolation between closely related species is often maintained by direct F1 hybrid sterility or hybrid breakdown in subsequent generations. This phenomenon is also observed in microorganisms within species of the *Saccharomyces* clade, including the model organism *Saccharomyces cerevisiae*. Members of *Saccharomyces* intermate and historical hybridization events have left genomic footprints, but these hybrids are themselves sterile and produce very few viable offspring. Previous work suggests a main cause of F1 hybrid sterility is the action of mismatch repair machinery. The sequence divergence between two parent *Saccharomyces* species—from ∼10% to ∼20%—is recognized by the cell as an aberrant attempt to recombine nonhomologous chromosomes during meiosis I, leading to massive nondisjunction and aneuploidy. Deletion of mismatch repair machinery improves fertility in hybrids of *S. cerevisiae* and its sister species *S. paradoxus* but leads to increased mutation accumulation during normal mitotic growth. To overcome this drawback, conditional alleles that specifically repress expression of mismatch repair machinery and the RecQ helicase *SGS1—*a major regulator of mitotic and meiotic recombination—during meiosis have been applied in *S. cerevisiae* x *S. paradoxus* hybrids with success. Here, we describe the application of the meiotic null system to a different hybrid: *S. cerevisiae* x *S. uvarum*. The system does not increase hybrid fertility likely due to increased sequence divergence and chromosomal translocations, but mass spore recovery and flow cytometry enable the recovery of near-haploid offspring which inherit an assortment of parental chromosomes. *S. uvarum* chromosomes which have undergone translocations since *S. cerevisiae* divergence were found to be co-inherited in viable progeny. Strikingly, these viable near-haploid hybrid offspring predominantly inherited Chr XII—carrying the critical rDNA repeat array—from the *S. uvarum* parent. These populations of mixed hybrid genomes can be leveraged in phenotypic studies to further understand the effects of hybrid genomes on organismal fitness.

## Introduction

### History of *Saccharomyces* hybrids

A powerful feature of *Saccharomyces cerevisiae* as a model organism comes not just from the species itself, but also from its relationship to the other closely related species within the *Saccharomyces* clade^1–3^. Although pairwise sequence divergence between *S. cerevisiae* and the other clade members varies up to ∼20%, *S. cerevisiae* is capable of cross-species mating with other clade members, resulting in interspecific hybrids^4,5^. *Saccharomyces* hybrids exist both naturally and in domesticated settings where the pressure of natural selection is imposed by humans^6–8^. There is even evidence that the entire *Saccharomyces* clade began with a common ancestor resulting from hybridization, followed by whole genome duplication^9–12^. Hybridization between two different *Saccharomyces* species can be an evolutionary advantage: benefits can arise from the genetic diversity and new allelic combinations that result from the combination of two parental genomes^13–17^. Hybrids can also suffer from this genetic mixing, displaying incompatibilities and/or accelerated genomic changes vs. intraspecies mating due to increased instability of the hybrid genome^18–22^.

While *Saccharomyces* yeasts can form hybrids, these hybrids are typically sexually reproductive dead ends because they are largely infertile (Figure 1A.)^23,24^. Hybrid infertility is not an uncommon phenotype and can be found throughout the tree of life including in mammals, plants, and microorganisms such as budding yeasts^25–29^. This infertility is viewed as one of many mechanisms that play a role in speciation. The mechanisms of hybrid infertility vary within the taxonomy of the hybrid’s parents: in mammals Dobzhansky-Muller genomic incompatibilities commonly prevent most F1 hybrids from producing F2 offspring^30–33^. In contrast, the infertility of plant hybrids is usually caused by abnormal ploidy in the hybrid compared to the parents^34,35^. These observations were the result of decades of early study however, and the last few decades have revealed no firm rules regarding the mechanisms of speciation and hybrid infertility in the Kingdoms of life^36^.

**Figure 1.**
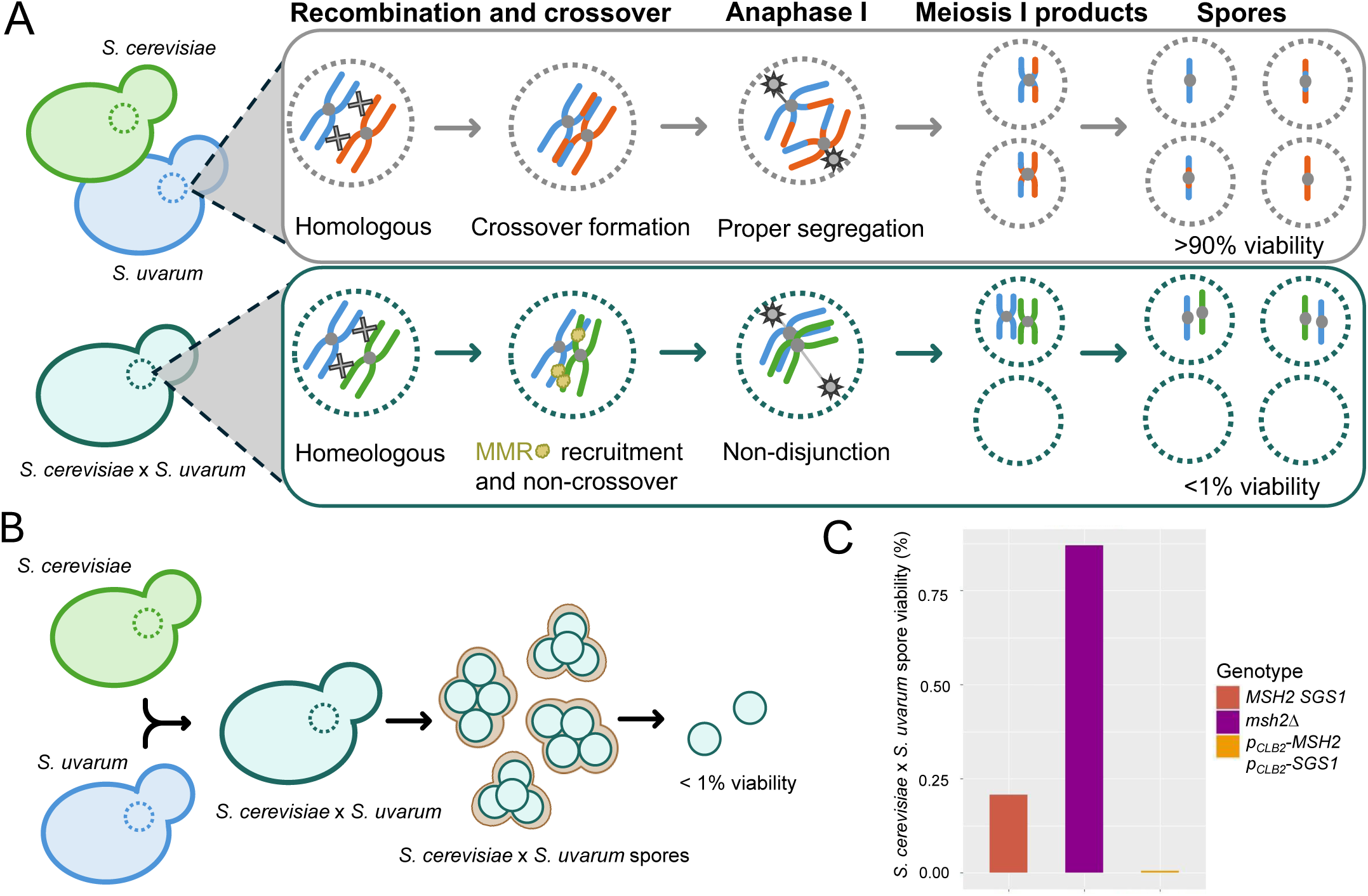
Depiction of *Saccharomyces* hybrid sterility and underlying molecular mechanism to target strategies for increasing viable segregants. [A] Diagram of representative chromosome pair comparing intraspecies Meiosis I mechanisms versus aberrant interspecies Meiosis I which results in aneuploidy and low offspring viability. [B] *Saccharomyces* species readily form hybrid diploids capable of sporulating but recovery of viable offspring is extremely low. [C] *S. cerevisiae* x *S. uvarum* hybrid viability results for a wild type, homozygous null allele of mismatch repair factor *MSH2*, and homozygous double meiotic null *MSH2* and DNA helicase *SGS1* alleles.

### Sterility of hybrids caused by anti-recombination activity of Mismatch Repair (MMR)

After determination of *Saccharomyces* hybrid spore inviability, the causality of this infertility became a topic of focus. There is some evidence that Dobzhansky-Muller genomic incompatibilities between parental genomes play some part in the infertility of certain hybrids, but it has been hypothesized that most hybrid infertility is due to the action of the mismatch repair (MMR) pathway during hybrid meiosis^37^. MMR machinery proteins are highly conserved across the tree of life and are critical to the cell for maintenance of genomic integrity^38–40^. MMR is most famously known for its role in repairing errors introduced during mitosis where it detects base - base mismatches and insertion/deletion mis-pairs generated after DNA replication and recombination. However, MMR also plays an important role during meiosis to ensure fidelitous chromosome segregation^41,42^.

MMR carries out this important role during sexual reproduction by ensuring efficacious recombination between homologous sequences during Prophase I of meiosis. The quality control provided by the pathway performs a critical function in detecting proper cross overs between parental homologous chromosomes. When there is a certain level of sequence divergence between the recombining parental genomes, MMR views the parental chromosomes as homeologous instead of homologous. The pathway then acts to abort the formation of any recombination intermediates by resolving the intermediates such that no crossover event will occur between the parental genomes (Figure 1B.)^43,44^.

Without recombination events among the parental genomes, there is not enough mechanical tension created by the physical connections of the chiasma between the homeologous chromosomes to properly line up on the metaphase plate before anaphase begins and chromosomes are distributed to daughter cells^41,45^. This mechanism is the current major hypothesis for the underlying molecular mechanism of hybrid infertility in *Saccharomyces*. Dysfunctional metaphase leading into anaphase causes massive aneuploidies in offspring via chromosome nondisjunction, rendering almost all inviable. The level of sequence divergence which serves to trigger MMR to block homeologous recombination is low enough that even intraspecies diploids of various *S. cerevisiae* strains display decreased spore viability. A sequence divergence of 1% is enough to observe a reduction of spore viability from 95% to approximately 80%^46^.

The lack of sexual reproduction within *Saccharomyces* hybrids is a major source of frustration for the field. The lack of sexual reproduction effectively precludes hybrids and their progeny from many of the classic genetic mapping strategies used within *Saccharomyces* species. Without the allelic shuffling and genetic diversity among offspring that is generated during meiosis, it becomes immensely difficult to study genotype to phenotype connections in hybrids. Classical approaches like Quantitative Trait Locus (QTL) mapping, which have been applied effectively for intraspecies *Saccharomyces* diploids, are rendered completely useless in the context of hybrid genetics. Overcoming the obstacle of *Saccharomyces* hybrid infertility is the focus of a considerable body of past and ongoing work.

### Loss of function and conditional MMR alleles to improve hybrid spore viability

A potential solution to the hybrid infertility issue was raised when mutations in the prokaryotic MMR factors *mutL*, *mutS*, and *mutH* were found to greatly increase the recombination rate between *Escherichia coli* (*E. coli*) and *Streptococcus pneumonia*—up to 1000X more over wild type MMR genes^47–50^. At the time, the homologs of *mutL* and *mutS* had been identified in *S. cerevisiae* as *PMS1* and *MSH2* respectively^51^. The functional conservation with their bacterial counterparts was suggested by several observations: both *PMS1* and *MSH2* mutants in yeast had an increase in the segregation frequency of genetic markers indicative of unrepaired heteroduplex DNA, similarly to the prokaryotic homologs^52–54^. In addition, both mutants exhibit a mutator phenotype further suggesting these strains also have poorer fidelity of DNA mismatch repair in mitotic growth as well.

All of these results together prompted Hunter et. al to investigate the effect of MMR mutations and deletions of pathway members on recombination rate and hybrid fertility in *Saccharomyces* yeasts^45,51^. The group did so by mating laboratory *S. cerevisiae* and *S. paradoxus* strains and analyzing spore viability after sporulation. At the time, *S. paradoxus* was identified as *S. cerevisiae*’s most closely related sister species, with the genomes estimated to have ∼10% divergence^53,55,56^. Of further note, karyotyping at the time had confirmed that there also existed no major structural rearrangements between the two species’ genomes, suggesting extensive synteny. These hybrid strains were then sporulated to induce meiosis via nutrient starvation and their tetrad ascospores manually dissected.

The rate of surviving progeny for each strain was compared to the spore viability of the wild type hybrid, which was ∼1%. The group found that by disrupting *MSH2* and *PMS1* individually, spore viability for the *S. cerevisiae* x *S. paradoxus* hybrid was increased to 10.2% and 7.2% respectively. In addition to this, and further supporting the important role of anti-recombination in hybrid infertility, surviving spores were found to have at least a two-fold increase in recombination events compared to wild type. These results laid the basis for continuing research that would attempt to further increase hybrid fertility.

Another more recent method that made a great stride in tackling this problem came from work by Duncan Greig’s group^57,58^. Building upon the previous work in MMR factors and anti-recombination genes more generally, Bozdag et al. targeted *MSH2* (the yeast MutS homolog) and *SGS1*, a DNA helicase previously found to also be involved in preventing homeologous recombination by rejecting aberrant recombination events, such as between non-homologous sequences which occur during hybrid meiosis, and instead resolving the DNA double stranded breaks as non-crossover events during meiosis I^59–61^.

As important as *msh2* mutants have been for studying the role of anti-recombination in hybrid inviability, it is critical to keep in mind that any MMR mutants exhibit a mutator phenotype. In vegetative growth these strains have an increased mutation accumulation rate. Previous work focused on reducing the mutation accumulation by limiting mitotic growth to only what was strictly necessary to gain enough cells to induce sporulation. While limiting periods of mitotic growth minimizes deleterious effects from random mutations, in an ideal world a trade off could be avoided between hybrid spore viability and mutation accumulation. One way to avoid the mutator phenotype is to turn anti-recombination genes off specifically only during meiosis so that their critical DNA repair functions during vegetative growth can be maintained. Leveraging this idea and the existence of previous meiotic null alleles, Bozdag et al. replaced the native promoters of *MSH2* and *SGS1* with the promoter sequence for the B-type cyclin gene *CLB2*. Clb2 plays an important role during the cell cycle by promoting the transition from G2 to M phase, after which the protein is specifically and efficiently marked for degradation and its expression silenced for proper cell cycle progression^62,63^. By placing both *SGS1* and *MSH2* under the *CLB2* promoter, their expression will specifically shut off only during meiosis.

Without the presence of *MSH2* and *SGS1*, it was hypothesized that homeologous recombination in the hybrids would proceed and result in proper chiasmata formation and crossing over leading to viable progeny from the lack of chromosome nondisjunction. This is indeed what Bozdag et al. found: by creating meiotic null alleles of *MSH2* and *SGS1* under the control of the *CLB2* promoter, *S. cerevisiae* x *S. paradoxus* hybrids spore viability increased up to 70-fold. This marked increase is almost matched to fertility rates of diverged non-hybrid crosses, and euploid offspring recovered showed multiple recombination events between each set of homeologous parental chromosomes.

The drastically increased fertility and the ability to recover euploid offspring with successful genomic recombination makes it possible to perform genetic mapping in otherwise sterile hybrids. We aimed to test this meiotic null approach as a method to also enable genetic mapping in *S. cerevisiae* x *S. uvarum* sterile hybrids, which have historically been less well studied within the field of *Saccharomyces* hybrid fertility research.

## Results

### A *msh2Δ S. cerevisiae* x *S. uvarum* hybrid exhibits slightly increased spore viability

Previous work in MMR mutants related to hybrid fertility focused on *S. cerevisiae* x *S. paradoxus* hybrids due to lower sequence divergence and full genomic synteny. This work aimed to understand how MMR mutant alleles affect hybrid spore viability in *S. cerevisiae* x *S. uvarum* hybrids with increased sequence divergence and translocations. We first tested the consequences of *MSH2* deletions on *S. cerevisiae* x *S. uvarum* hybrid spore viability (Figure 1C.). We sporulated a hybrid diploid generated from wild type haploids from each species and manually dissected and scored progeny. After growth, percent viability was calculated to be ∼0.87%. This value is very similar to previously reported data of ∼1% for other similarly diverged inter-species *Saccharomyces* hybrids.

After repeating this experiment on the *msh2Δ*/*msh2Δ* hybrid, we observed this percent viability increased to around 2.4%. To test spore viability for meiotic null *MSH2* and *SGS1* in *S. cerevisiae* x *S. uvarum* hybrids, the native promoters of *MSH2* and *SGS1* were replaced by each species’ *CLB2* promoter. Previous comparative expression analysis was referenced to ensure conserved and specific repression of each species’ *CLB2* during meiosis^64^. The viability of these offspring showed nowhere near the increase seen for the engineered *p_CLB2_ S. cerevisiae* x *S. paradoxus* spores: viability was much lower than for the *msh2Δ* hybrid strain progeny.

These numbers are much smaller than the ∼10% previously observed for *S. cerevisiae* x *S. paradoxus* hybrids, which share more sequence identity and possess more syntenic genomes. Reducing the action of anti-recombination increases homeologous recombination in *S. cerevisiae* x *S. paradoxus* hybrids but there exist additional barriers to hybrid fertility in *S. cerevisiae* x *S. uvarum* hybrids. Larger chromosome scale translocations have been documented between the two species’ genomes, providing an additional known barrier to hybrid fertility. However, chromosomal translocations alone cannot explain the lower hybrid fertility found in *S. cerevisiae* x *S. uvarum* hybrids so other potential factors must also contribute^58^.

Meiotic null MMR *S. cerevisiae* x *S. uvarum* hybrids remain infertile but offspring aneuploidy is decreased.

In order to further assess the genomic composition of surviving progeny, an en masse spore retrieval protocol known as random spore analysis (RSA) was utilized (Figure 2). RSA is facilitated by using a parent *S. cerevisiae* strain engineered to contain a suite of markers known as the Synthetic Genetic Array (SGA). These markers enable efficacious bulk selection of haploid *MATa* progeny from a sporulated diploid. We utilized the haploid selection features but omitted the mating type specific selection to maximize number of recovered hybrid offspring. Offspring recovery occurs by subjecting the hybrid parent sporulation culture to a combination of mechanical and chemical selections to selectively kill diploids and mechanically separate spores. A key element of the SGA marker suite for our purpose was the *CAN1* gene, which encodes for an arginine permease. A single functional copy of *CAN1* facilitates the import of arginine or the lethal arginine analog canavanine into the cell, with the latter leading to cell death. Haploid offspring of a heterozygous diploid should only have one copy of the gene, thus progeny that inherit only the *can1* null allele survive while *CAN1* progeny and heterozygous diploids die. Leveraging RSA in this way, hybrid offspring for each experimental and control strain were collected and subjected to growth assays and DNA content staining followed by flow cytometry (Figure 2). Analysis was performed in the FlowJo software to compare histograms of relative fluorescence units for asynchronously growing cells collected for each hybrid segregant.

**Figure 2.**
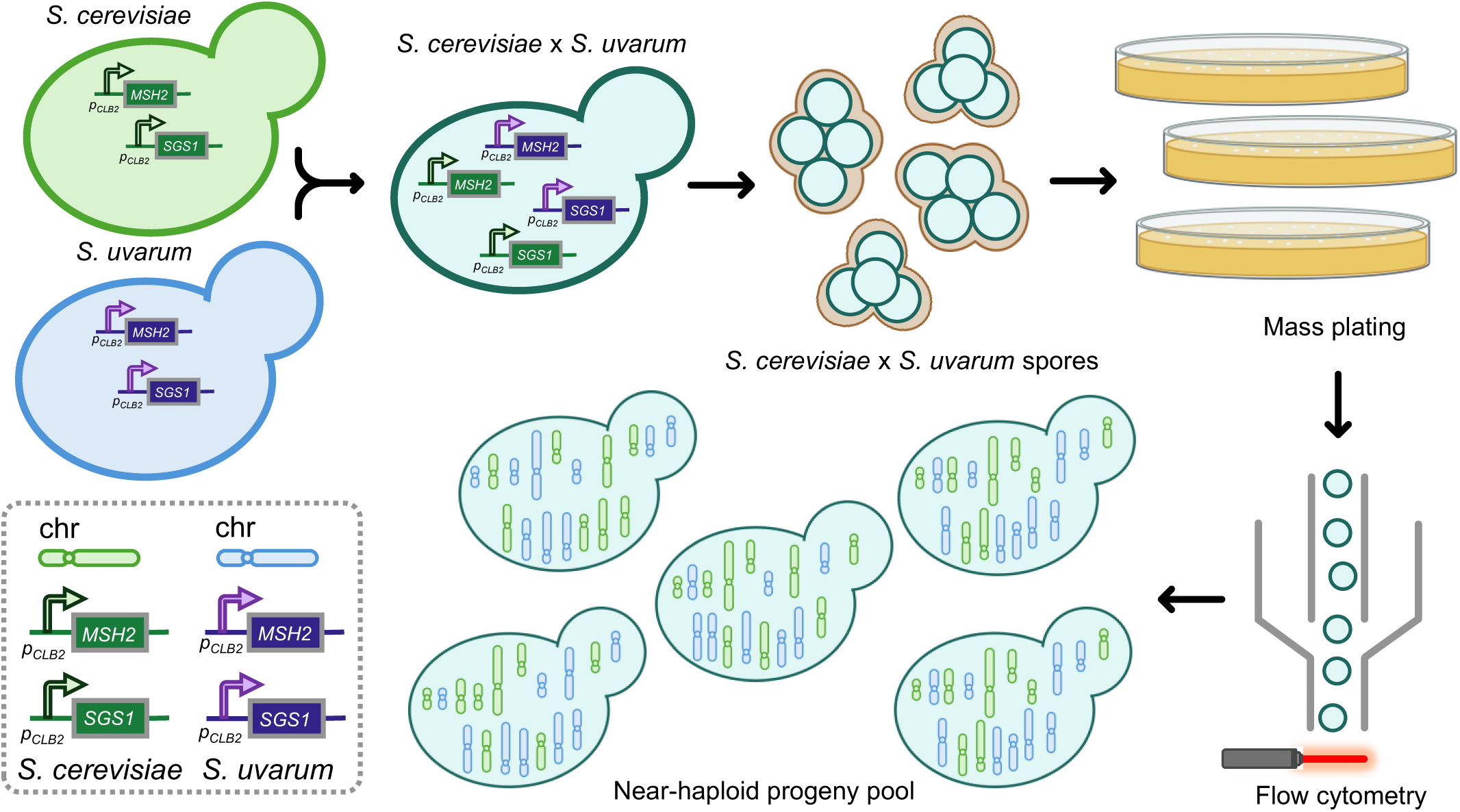
Diagram depicting workflow with constructed meiotic null *S. cerevisiae* x *S. uvarum* hybrid strain undergoing sporulation, recovery of surviving offspring with mass plating method, and flow cytometry analysis of chromosome content to identify near-haploid collection of progeny for further screening and phenotyping.

Specifically, these experimental segregant DNA content staining plots were compared by FlowJo fluorescence gating pairwise with the results for both a haploid and diploid control and cut offs were determined for labeling as near-haploid, aneuploid slight, or aneuploid severe (Figure 3A, Figure 3B.). The segregants were then binned into several different categories based on the overlap, or lack thereof, of the experimental samples with the haploid control (Table 1). These flow cytometry results were used to calculate a percentage of near-haploid offspring for each hybrid population: wild type, and two different biological replicates of the homozygous *p_CLB2-MSH2_* and *p_CLB2-SGS1_* engineered strain (Figure 3C.). As expected, the wild type hybrid strain had the most aneuploid offspring, with the near-haploid percent at only 7.89%. The biological replicates for the experimental *p_CLB2_* promoter replacement hybrids yielded a substantially larger percentage of near-haploid offspring at 46.5% and 43.9% respectively. In addition, the aneuploid offspring produced by these *p_CLB2_* replacement strains were on average less aneuploid, indicated by the progeny classified as slight aneuploid vs severe.

**Figure 3.**
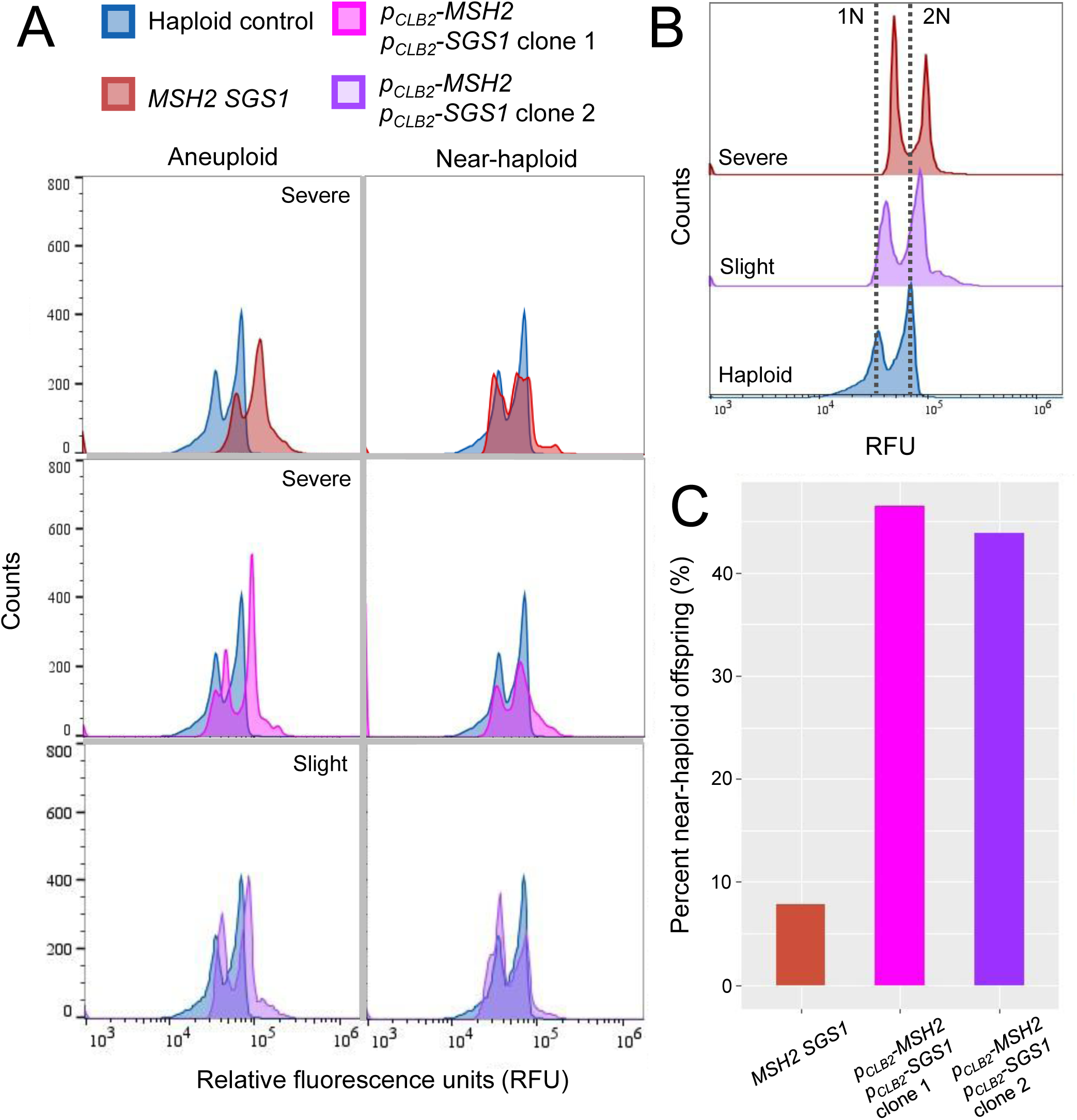
Comparative DNA staining flow cytometry analysis of hybrid offspring ploidy between wild type and *p_CLB2_-MSH2* p*_CLB2_-SGS1* meiotic null strains. [A] Representative plots are shown for binning of *p_CLB2_* progeny as Severe or Slightly aneuploid. [B] Offset overlay showing in more detail the average difference in aneuploidy levels for *p_CLB2_* offspring vs. wild type offspring. [C] Comparison of chromosome content for progeny of diploid hybrid strains. All strains are homozygous at all indicated alleles. Total number of offspring analyzed: 38 *MSH2 SGS1*, 43 *p_CLB2_-MSH2* p*_CLB2_-SGS1* clone 1, 41 *p_CLB2_-MSH2* p*_CLB2_-SGS1* clone 2.

**Table 1.** Flow cytometry ploidy binning summary for recovered progeny of wild type and *MSH2 SGS1* meiotic null *S. cerevisiae* x *S. uvarum* hybrids.

| MMR Genotype | Severe aneuploid | Slight aneuploid | Near-haploid | % near-haploid |
| --- | --- | --- | --- | --- |
| *MSH2 SGS1* | 35 | 0 | 3 | 7.89 |
| *pCLB2-MSH2 pCLB2-SGS1*<br>Clone 1 | 9 | 14 | 20 | 46.5 |
| *pCLB2-MSH2 pCLB2-SGS1*<br>Clone 2 | 6 | 17 | 18 | 43.9 |

### Whole genome sequencing of select progeny reveal patterns of chromosome inheritance

Based on the results of the flow cytometry DNA content screens, we selected 16 *p_CLB2_-MSH2 p_CLB2_-SGS1* hybrid strain progeny that appeared near-haploid for 10X low coverage whole genome sequencing (WGS). The low coverage sequencing was sufficient to properly map reads and determine relative copy number between the two parental genomes, given the 20% sequence divergence between them. Analyzing the alignment of these reads to a hybrid parent genome revealed that almost all the progeny had inherited a complete set of 16 chromosomes with limited aneuploidies. Interestingly, we did not see uniparental inheritance: instead each progeny inherited each chromosome mostly in its entirety from either the *S. cerevisiae* or *S. uvarum* parent or both in the cases of aneuploidies (Figure 4A.). A summary of gross chromosome content for the recovered viable progeny was as follows: four with 17 chromosomes, six with 18 chromosomes, five with 19 chromosomes, and one with 21 chromosomes.

**Figure 4.**
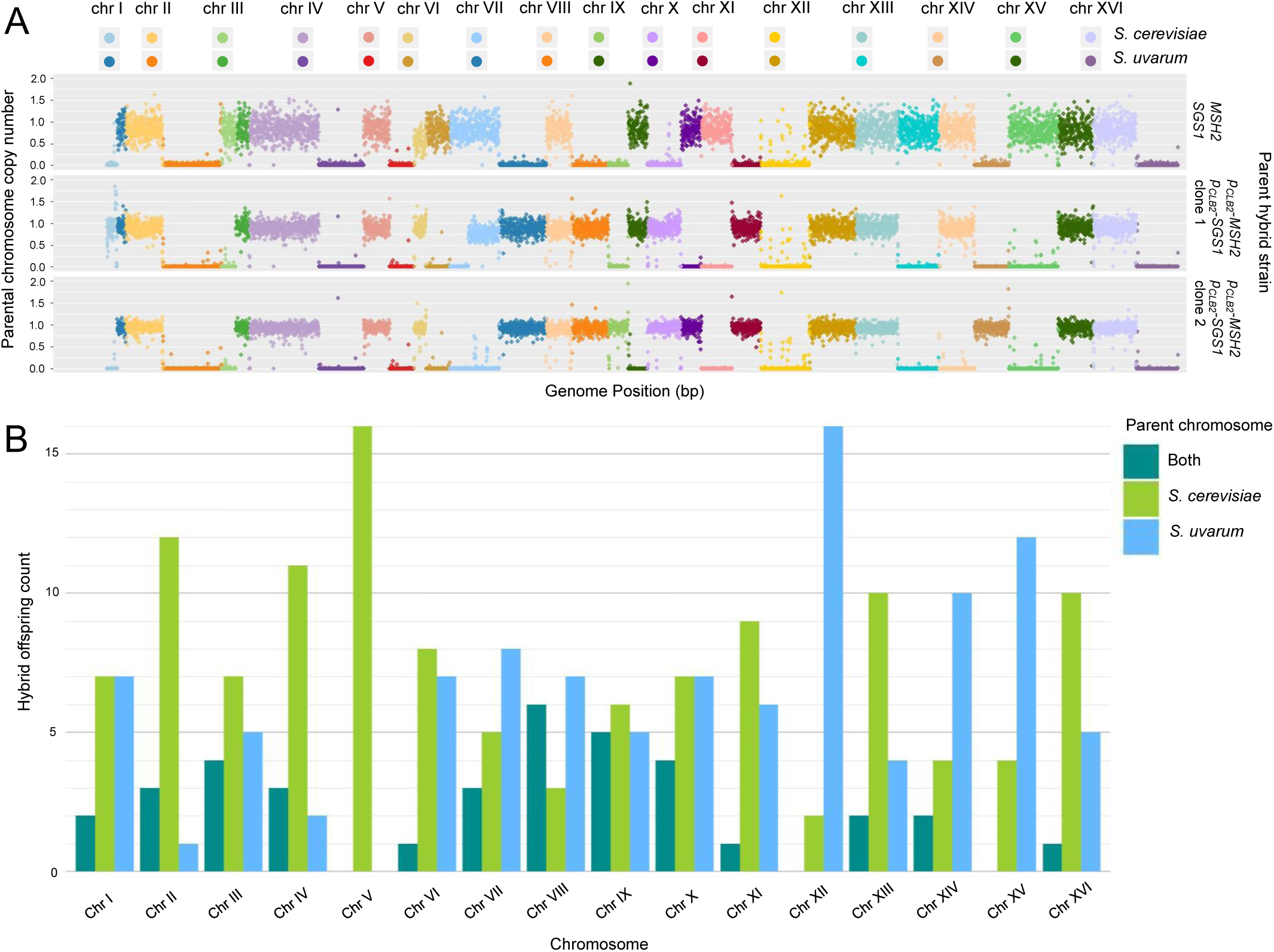
Summary view of parental chromosome inheritance for viable *S. cerevisiae* x *S. uvarum* offspring. [A] Copy number variation plots from alignment of WGS reads to concatenated *S. cerevisiae* and *S. uvarum* hybrid genome. Representative offspring from wild type and either engineered clonal isolate showing patterns of coinheritance and a recombination event. Data for all offspring can be found in Supplementary file 1. [B] Summary plot of parental chromosome inheritance for the 16 viable meiotic null offspring recovered. and double inheritance like Chr X. Selecting for uniparental inheritance of these chromosomes could be used in future studies to inform the mechanism of coinheritance of certain aneuploidies.

Recombination between the hybrid genomes was rare, observed in only a single recovered near-haploid offspring from a meiotic null parent. Particular attention was directed to the inheritance patterns for chromosomes that contain one of the translocations present between the *S. cerevisiae* and *S. uvarum* genomes: reciprocal for Chr II and Chr IV, Chr VIII and Chr XV, Chr VI and Chr X, and nonreciprocal for left arm of Chr VII onto Chr V. The presence of homeologous sequence tracts between these translocated parental chromosomes suggests coinheritance for each pair across the 16 near-haploid progeny reported here. Viable hybrid progeny inherited both of one chromosome and a single chromosome from one parent, or inherited parent matched chromosome pairs: for example both Chr II and Chr IV from *S. cerevisiae* only (Figure 4B.). Based on previous studies in fungi for the consequences of translocations during meiosis, this finding aligns with the expectation that during meiosis chromosomes with homeologous translocated sequences will form quadrivalents on the metaphase plate and could thus cosegregate during anaphase I^65^.

Other striking patterns observed across the 16 near-haploid progeny (Supplementary file 1) were observed for parental chromosome inheritance (Figure 4). One example is that all progeny recovered had only one copy of Chr V, and it was always inherited from the *S. cerevisiae* parental genome. This pattern is expected due to the use of canavanine to select for the *can1* allele from *S. cerevisiae,* as described above. This provides a method to bias parental chromosome inheritance from meiotic null *S. cerevisiae* x *S. uvarum* parents. Usage of selectable and counterselectable markers could also be used to select for single chromosome inheritance for chromosomes that show enriched aneuploidy

Another example of strong parental bias for chromosome inheritance was observed for Chr XII, where over 85% of viable *S. cerevisiae* x *S. uvarum* offspring had inherited only the *S. uvarum* chromosome XII, and no progeny inherited both parental Chr XII. This finding is particularly intriguing due to the presence of the multicopy rDNA array on the right arm of Chr XII for both *S. cerevisiae* and *S. uvarum*. The rDNA exists as a tandem array of multiple copies, with an observed range of approximately 100-200 copies in most *S. cerevisiae* isolates. Each single copy encodes for the RNA transcript components of the cell’s ribosomes: the nucleoprotein complex responsible for translation of the proteome from RNA transcripts. Previously, work on understanding the phenotypic consequences of rDNA variation and its role in the cell’s biological functions had focused almost exclusively on building this understanding within intraspecies variation among *S. cerevisiae* isolates with little consideration of rDNA phenotypic effects in *Saccharomyces* hybrids. While the viability of the parent *S. cerevisiae* x *S. uvarum* hybrid suggests no fundamental incompatibility between *S. cerevisiae* or *S. uvarum* rDNA and the rest of the hybrid genome, it is possible that the process of undergoing meiosis provides a selection pressure which reveals an incompatibility resulting in biased inheritance of the *S. uvarum* Chr XII in viable hybrid progeny. The strong signature of *S. uvarum* Chr XII inheritance bias, even for a small population, is striking and points at potential biological significance.

Assessing aneuploidy of parental chromosome bias across the collection of near-haploid genomes reveals suggested inheritance patterns as well. We observed an enrichment (almost a third of recovered progeny) in biparental aneuploidy for chromosomes VIII and IX. For uniparental inheritance bias (defined here as greater than 2:1 ratio of parental inheritance per chromosome with minimal aneuploidy), we observed a bias for *S. cerevisiae* parental inheritance for chromosomes II, IV, V, XIII, and XVI. An enrichment for *S. uvarum* parental inheritance was observed for chromosomes XII, as noted above, XIV, and XV. Other chromosomes were observed to be inherited equally from either parental genome, including chromosomes I, III, V, VII, X, and XI. We do not currently understand the reasons for these patterns or yet have an understanding of the instances of genetic conflict that could potentially be occurring. These represent research areas of interest to be further pursued with this engineered hybrid system.

## Discussion

The work presented here examined the effect of mutant alleles for mismatch repair factors on hybrid spore viability for interspecies *S. cerevisiae* x *S. uvarum* hybrids. Previous work has focused solely on assessing the spore viability effect of suppressing the anti-recombination activity of MMR in hybrids of *S. cerevisiae* and its sister species *S. paradoxus*^66^. Meiotic null alleles of the MMR factor *MSH2* and RecQ helicase *SGS1* were previously observed to increase spore viability of *S. cerevisiae* x *S. paradoxus* from less than 1% to approximately 35% with multiple recombination events in the hybrid genome during meiosis. Here we report that replicating the meiotic null alleles in *S. cerevisiae* x *S. uvarum* hybrids revealed no increase in fertility. While disappointing for efforts to genetically map traits for hybrids, the stark difference in spore viability for *S. cerevisiae* x *S. paradoxus* versus *S. cerevisiae* x *S. uvarum* hybrids is not entirely surprising. In *S. cerevisiae* x *S. paradoxus* hybrids, the parental genomes have less sequence divergence at ∼10% (∼20% for *S. cerevisiae* vs *S. uvarum*) and are more syntenic (3 reciprocal and 1 nonreciprocal translocations for *S. cerevisiae* vs *S. uvarum*)^67^.

Previous studies that have examined the effect of translocations on hybrid sterility and more recent work in intra-species matings of lab and natural *S. cerevisiae* isolates have shown that translocations have a separate and significant impact on hybrid fertility from genomic sequence divergence. Further supporting this hypothesis is a study where the authors found that in a screen of intra-species *S. cerevisiae* crosses between the lab S288C reference strain and natural isolates the subsequent F1 hybrids with the lowest fertility all contained translocations in comparison to the *S. cerevisiae* reference genome^68^. This intra-species crossing screen also revealed that progeny viability was not correlated with sequence divergence for these intra-species hybrids, but instead found reproductive isolation was mainly related to genomic rearrangements. In this scenario, the effect of sequence divergence between parental genomes is minimized and it becomes clear that in the case of low sequence divergence genomes, translocations alone do contribute to driving hybrid infertility.

We believe that a similar effect could be at play in *S. cerevisiae* x *S. uvarum* hybrid fertility, and could potentially suggest a partial explanation for the lack of offspring even after the *p_CLB2_* MMR meiotic null alleles: by alleviating the action of mismatch repair as a cause of hybrid infertility, the impact of multiple reciprocal and nonreciprocal translocations remain and still severely impact hybrid meiosis in a hybrid with greater sequence divergence (20%) then previously examined for the effect of translocations between hybrid genomes on spore viability^69^. These previous results and the work reported here suggest that the effects of translocations and sequence divergence between two *Saccharomyces* genomes can be overlapping and independent in their severity of effect for hybrid spore viability. However, this explanation still does not fully account for the complete lack of increased hybrid spore viability in the *S. cerevisiae* x *S. uvarum* meiotic null backgrounds. One approach to detangle the effect of translocations vs sequence divergence on *S. cerevisiae x S. uvarum* spore viability would be to genetically engineer *S. uvarum* strains with reversed translocations to reestablish genomic synteny with *S. cerevisiae*: a method previously utilized to study hybrid sterility in *S. cerevisiae* x *S. mikatae* hybrids^67^.

The lack of any recovered viable progeny from manual dissections of *p_CLB2-MSH2_ p_CLB2-SGS1_ S. cerevisiae* x *S. uvarum* hybrids is striking. The molecular mechanism for this lack of progeny when compared to *MSH2 SGS1* hybrids is unknown. It could be speculated that due to the critical action of Sgs1 in resolving meiotic recombination intermediates to non-crossovers for nonhomologous sequences, the loss of this function during *S. cerevisiae* x *S. uvarum* hybrids has particularly detrimental consequences due to the additional problematic nature of translocations and increased sequence divergence between the parent genomes when compared to the *p_CLB2_-SGS1* allele in *S. cerevisiae* x *S. paradoxus* hybrids.

Another approach in future pursuits would be to dissect patterns of viability and percent near-haploid offspring for both *p_CLB2_-MSH2* and *p_CLB2_-SGS1* single meiotic null hybrid strains. These single meiotic null hybrids will enable interrogation of the underlying molecular mechanisms for the total lack of recovered viable offspring during manual dissection by separating potential effects of meiotic null *SGS1* from *MSH2*.

Future work to increase hybrid spore viability could also focus on other actors during Meiosis I prophase that play a role in preventing or increasing homeologous recombination, including the DNA repair protein Rad51 and its meiotic specific partner Dmc1, as well as important members of the synaptonemal complex—a proteinaceous assembly that forms between homologous chromosomes in Prophase I and plays a critical role in regulating crossover formation^70^. Additionally, the return-to-growth approach which involves prematurely aborting meiosis after recombination was previously shown to increase recovery of hybrids that have recombined genomes^71^. It is also possible to dodge the challenges of hybrid meiosis by creating tetraploid hybrids where homeologous recombination between parental strains does not occur but allelic shuffling can still occur internal to each parental genome and its phenotypic consequences in the hybrid studied by QTL^72^. Other studies have suggested an important role in environmental conditions for *S. cerevisiae* and *S. uvarum* meiotic progression timing and recombination rates, offering another variable to explore for improving meiotic success and recombination in hybrids and increasing recovery of viable progeny^73^. Understanding the deeper reasons for lack of fertility increase in these engineered *S. cerevisiae* x *S. uvarum* hybrids could also potentially lead to other approaches that will provide an increase in fertility.

While the engineered promoter replacement *S. cerevisiae* x *S. uvarum* hybrids may not have yielded an increased percentage of viable spores by manual dissection, the use of higher throughput random spore analysis and flow cytometry shows that spores recovered from the engineered strain show a significant increase in near-haploid offspring compared to the progeny generated from the *MSH2 SGS1* wild type hybrid strain. It should be noted that the particular random spore method applied here yielded a significant proportion of diploid parental strain yeast which survived the vegetative cell killing treatment. While a more stringent spore selective protocol can be utilized to overcome this drawback, it is also possible to sort away this population by flow cytometry while selecting for near-haploid offspring.

A serious limitation of this study was the number of near-haploid progeny studied. While none of the progeny recovered were perfectly haploid, there exists the possibility that with a higher number of examined offspring truly haploid offspring with mixed chromosomal inheritance could be recovered and subjected to further study. With the meiotic null *S. cerevisiae* x *S. uvarum* hybrids it is also possible to bias parental chromosome inheritance and select against biparental inheritance, as evidenced by the *S. cerevisiae* Chr V unilateral inheritance due to the usage of the *CAN1* gene as a selectable marker during the mass spore recovery protocol. Similarly, use of the SGA mating type-specific selection markers could bias Chr III inheritance, and also generate populations consisting of a single mating type, facilitating pooled experiments.

Meiotic null engineered hybrids in tandem with mass plating methods and flow cytometry sorting, instead of the non-sorting flow cytometry utilized in this study, empower recovery of a population of viable near-haploid progeny that have mixed inheritance of *S. cerevisiae* and *S. uvarum* chromosomes (Figure 2.). Individual isolates or the pooled library of mixed chromosome near-haploid offspring can then be screened or phenotyped to study trans acting genetic background effects and the genomic consequences of mixed parental chromosomal inheritance in hybrid offspring in short or long experimental timeframes as has been done for *S. cerevisiae* x *S. paradoxus* hybrid progeny^74^. In conclusion, we present here a method by which MMR meiotic null *S. cerevisiae* x *S. uvarum* hybrids can be sporulated and their surviving near-haploid progeny can be isolated through mass plating methods and flow cytometry to enable the application of genetic mapping methods to infertile *Saccharomyces* hybrids separated by genomic translocations.

## Materials and methods

### Strains

Detailed strain genotype information found in Supplemental Table 1.

For the *msh2* knockout strains, the *S. cerevisiae* haploid was sourced from the *MATa* Yeast Deletion Collection. The creation of the SGA version of this collection was described previously in Tong et al.^77^, but in short, each non-essential gene was systematically deleted by a barcoded KanMX cassette within a diploid strain derived from BY4742 already containing an assortment of other markers. Subsequent sporulation of this diploid and selective screening of progeny allowed for recovery of haploid offspring with the SGA marker suite, and the gene of interest deletion cassette. The *msh2Δ* knock out for *S. uvarum* was generated previously. Briefly, the endogenous genomic locus of *MSH2* was replaced by transforming a wild type *S. uvarum* yeast strain derived from CBS7001 with a PCR product containing the *NatMX* selectable marker flanked by primer designed homology to the endogenous *MSH2* locus.

*S. cerevisiae* x *S. uvarum* hybrids were generated by patch mating of parental strains on solid YPD plates and picking zygotes under the microscope after 4-5 hours post mating. A ‘wild type’ hybrid was generated by mating a haploid version of the SGA strains termed YMD895 to the haploid *S. uvarum* strain termed YTW2.

The haploid *MSH2* and *SGS1 S. cerevisiae* strains used in this work were generated using a diploid *S. cerevisiae* parental strain originating from the synthetic genetic array (SGA) collection originally generated by Tong et al^77^. This strain had the genotype of *can1Δ::STE2-Sp_his5*/*CAN1 lyp1Δ*/*LYP1 his3Δ1*/*his3Δ1 leu2Δ0*/*leu2Δ0 ura3Δ0/ura3Δ0 met5Δ0/ LYS2/LYS2 MET15/met15Δ0* and was made from BY4742, which is isogenic to the reference laboratory *S. cerevisiae* strain S288C. A diploid strain termed YTW3 was used to avoid any suppressor mutations in case of fitness defect in the promoter replacement allele that could occur in a haploid. Each promoter replacement strain was made by transforming the yeast with the appropriate PCR product (plasmid assembly defined below) as described previously^75^. Briefly, a colony of the parent *S. cerevisiae* strain YTW3 was grown overnight. This culture was diluted the next morning and grown until an OD of 0.5 – 0.8 was reached. Cells were then spun down, washed, and subjected to a chemical transformation using lithium acetate, polyethylene glycol, water, and DNA to be transformed. Cells were then recovered for two hours in YPD and finally plated on either D + G418, or D + CloNAT plates as needed. Single colonies were picked to check for promoter replacement via PCR and Sanger sequencing.

*S. uvarum* strains in this study all originated from the CBS7001 parental strain, whose HO locus has been disrupted to prevent mating type switching. Promoter replacement strains were made similarly to the *S. cerevisiae* strain: transformation of the diploid termed YTW8 with PCR products to replace either the *MSH2* or *SGS1* promoter with the *S. uvarum CLB2* promoter marked with a *NatMX* and *KanMX* cassette respectively. The only change in protocol was for the heat shock to occur at 37°C instead of 42°C, and for all normal growth to occur at 25°C instead of 30°C due to the difference in preferred growth temperature for *S. uvarum* compared to *S. cerevisiae*.

### Gibson assembly

To replace the native promoters of *SGS1* and *MSH2*, primers were designed to amplify the *CLB2* promoter sequence from genomic DNA of both *S. cerevisiae* and *S. uvarum*. These PCR products were then cloned into the plasmid pFA6a via Gibson assembly using inverse PCR for the plasmid backbone to remove several expression cassettes unnecessary for this experiment and designed overhangs. For each species’ promoter, two constructs were created: one with a *KanMX* cassette upstream of the promoter, and the other with a *NatMX* cassette. The *KanMX* cassette was used to demarcate the replacement of the native *SGS1* promoter with *p_CLB2_*, and the *NatMX* marker used to indicate promoter replacement for *MSH2* loci in both species.

### Spore viability

Hybrid cultures were sporulated in 5 mLs of 1% potassium acetate, 0.05% dextrose, and 0.1% yeast extract for 4-7 days. Ascospore dissection was carried out by spinning down 50 μL of the sporulation culture, resuspending in 15 μL of yeast lytic enzyme (YLE) at 100 U/mL in 1 M sorbitol, and plating on YPD after a 30 minute digestion at 30°C.

Dissections were carried out at room temperature on a Nikon H550S Eclipse 50i scope via manual needle dissection. After 3 days of growth at 25°C surviving tetrad spores were counted. Percent viability for each hybrid strain was calculated by the following formula: 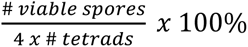.

To recover viable progeny en masse in a high throughput setting, the method of random spore analysis was used^76^. Briefly, a volume of sporulation culture for each hybrid was spun down and resuspended in 200 μL of 100 U/mL YLE. After 30 minutes of incubation at 30°C, cultures were subjected to 1 minute of vortexing followed by plating onto C +Canavanine -arginine -serine.

Surviving colonies were collected, preserved at -80°C in glycerol stocks, and subsequently subjected to growth analyses and flow cytometry

### Flow cytometry

DNA content flow cytometry was performed using a BD Accuri C6 flow cytometer. The protocol is as described previously in the 2015 edition of Methods in Yeast Genetics and Genomics (A Cold Spring Harbor Laboratory Course Manual)^78^. Briefly, spores as well as haploid and diploid controls were seeded to grow overnight at the appropriate temperatures. Cultures were back diluted to ∼0.2 OD the next day and then harvested between ∼ 0.5 – 0.8 OD by spinning down 1 mL and resuspending in 1 mL of 70% ethanol. Half of these fixed cells were then washed with water and subjected to RNase A digestion in 200 μL of 50 mM Tris-HCl (pH = 8.0) for 2 hours at 37°C. Samples were then spun down and resuspended with 2 mg/mL Proteinase K in 200 μL of 50 mM Tris-HCl (pH = 7.5) for 30 minutes at 50°C. After another spin, samples were resuspended in 200 μL of FACS buffer (200 mM Tris-HCl pH = 7.5, 200 mM NaCl, and 78 mM MgCl_2_). Directly before samples were loaded onto the cytometer, 10 μL of cells were transferred to a 96-well plate containing 300 μL of SYBR green diluted 5000X in 50 mM Tris-HCl (pH = 7.5) and subjected to sonication. All samples were sonicated for 3 seconds at 10% power, resulting in a 5-10 Watt power output. Analysis of flow cytometry samples, including gating for debris and doublets, was carried out in FlowJo.

### WGS Sequencing Preparation and Analysis

The sequencing libraries for the near-haploid hybrid offspring were prepared using a commercial Illumina NextSeq library sequencing kit. The quality and size of the libraries was verified by performing gel electrophoresis on a 5% SDS-PAGE gel for 40 minutes at 150 V followed by staining and imaging. An average 10X sequencing depth was achieved across each *S. cerevisiae* x *S. uvarum* near-haploid offspring genome. Final reads were processed as follows: Trimmomatic was used to remove any remaining adapter sequences, alignment of trimmed reads to a concatenated *S. cerevisiae* x *S. uvarum* genome was carried out by the Burroughs-Wheeler Aligner package (BWA-MEM), sorted using PicardTools, multimapping reads removed by a custom script (https://github.com/taykatwang/hybrid-meiotic-null), and the samtools package was then used to generate read depth files for each strain. Final copy number plots for all hybrid progeny can be found in Supplemental file 1.

### Sequencing Analysis Pipeline for Copy Number Variation

Read depth files from the initial analysis detailed above were then used as input to a custom copy number variation script written in RStudio (https://github.com/taykatwang/hybrid-meiotic-null). In brief, alignment files were imported to RStudio as data frames while simultaneously removing mitochondrial DNA and read depth at each base pair position was normalized to average genome wide read depth. For clarity of data visualization, normalized read depth was then condensed into 3 kb bins and per bin average read depth calculated. The extreme terminal ends of each chromosome were trimmed to remove the excess noise of read depth resulting from the low coverage. A custom script (https://github.com/taykatwang/hybrid-meiotic-null) was used to generate the x axis of entire concatenated *S. cerevisiae* x *S. uvarum* parental genome by manually retrieving per chromosome size for each species, and genome position added for each 3 kb bin to be plotted. Final plots were then generated for each near-haploid progeny that plotted copy number for each individual parental chromosomes along an x axis of the concatenated parental hybrid genome.

## Data Availability Statement

The data underlying this article are available in the NCBI Sequence Read Archive database and can be accessed with BioProject ID PRJNA1552138.

## Acknowledgements

We gratefully thank Dr. Duncan Greig and Dr. G. Ozan Bozdag for sharing pCLB2 meiotic null S. cerevisiae strains with the lab, and for graciously answering strain engineering and design questions for generating the alleles in our strain backgrounds. We thank Noah Fredstrom and Dr. Joe Armstrong for assistance with Illumina sequencing. We thank Dr. Monica Sanchez for being part of paving the way of *S. uvarum* work in the lab and for generating the *S. uvarum msh2Δ* strain. We thank Genome Sciences IT, including Charles Winston and Skylar Thompson, for support with retrieving and accessing packages required for sequencing analysis. We also thank the yeast media center in Genome Sciences run by the wonderful Sandra Pennington for all her support and technical media help. We would also like to thank Dr. David Gresham and Dr. Julie Chuong for helpful, constructive conversations during construction of the sequencing pipeline. We acknowledge that Artificial Intelligence was used at the end of R script development via ChatGPT 3.5 to assist in data visualization by contributing several lines of code supervised by the first author to arrange chromosome numbers in order on plot and in annotation of *S. uvarum* chromosomes. Finally, we would also like to extend our gratitude to Dr. Gavin Sherlock and Dr. Sasha Levy for stimulating discussion, suggestions, and support during this project.

## Funding

This work was supported by National Science Foundation grant 1516330 and by R01 HG010378 from the National Human Genome Research Institute at the National Institutes of Health.

MJD holds the William H. Gates III Endowed Chair in Biomedical Science.

## Notes

### Competing Interest Statement

The authors have declared no competing interest.

### Summary of Updates

New references added, some figure changes for increased clarity, and discussion expanded.

## References

1. Taylor, J. W. et al. Phylogenetic Species Recognition and Species Concepts in Fungi. Fungal Genet. Biol. 31, 21–32 (2000).

2. Naumov, G. I., James, S. A., Naumova, E. S., Louis, E. J. & Roberts, I. N. Three new species in the *Saccharomyces sensu stricto* complex: *Saccharomyces cariocanus*, *Saccharomyces kudriavzevii* and *Saccharomyces mikatae*. Int. J. Syst. Evol. Microbiol. 50 **Pt** **5**, 1931–1942 (2000).

3. Guillamón, J. M., Barrio, E., Huerta, T. & Querol, A. Rapid characterization of four species of the *Saccharomyces sensu stricto* complex according to mitochondrial DNA patterns. Int. J. Syst. Bacteriol. 44, 708–714 (1994).

4. Liti, G., Barton, D. B. H. & Louis, E. J. Sequence diversity, reproductive isolation and species concepts in *Saccharomyces*. Genetics 174, 839–850 (2006).

5. Dunn, B. & Sherlock, G. Reconstruction of the genome origins and evolution of the hybrid lager yeast *Saccharomyces pastorianus*. Genome Res. 18, 1610–1623 (2008).

6. González, S. S., Barrio, E., Gafner, J. & Querol, A. Natural hybrids from *Saccharomyces cerevisiae*, *Saccharomyces bayanus* and *Saccharomyces kudriavzevii* in wine fermentations. FEMS Yeast Res. 6, 1221–1234 (2006).

7. Sipiczki, M. Interspecies hybridization and recombination in *Saccharomyces* wine yeasts. FEMS Yeast Res. 8, 996–1007 (2008).

8. Barbosa, R. et al. Evidence of Natural Hybridization in Brazilian Wild Lineages of *Saccharomyces cerevisiae*. Genome Biol. Evol. 8, 317–329 (2016).

9. Kellis, M., Birren, B. W. & Lander, E. S. Proof and evolutionary analysis of ancient genome duplication in the yeast *Saccharomyces cerevisiae*. Nature 428, 617–624 (2004).

10. Wolfe, K. H. Origin of the Yeast Whole-Genome Duplication. PLoS Biol. 13, e1002221 (2015).

11. Wolfe, K. H. & Shields, D. C. Molecular evidence for an ancient duplication of the entire yeast genome. Nature 387, 708–713 (1997).

12. Adaptive genome duplication affects patterns of molecular evolution in *Saccharomyces cerevisiae* | PLOS Genetics. https://journals.plos.org/plosgenetics/article?id=10.1371/journal.pgen.1007396.

13. Haberkorn, C. et al. Adaptive benefits of hybridization in *Saccharomyces* yeast are constrained by genomic background and depend on temperature. Evolution 80, 1837–1855 (2026).

14. Stelkens, R. & Bendixsen, D. P. The evolutionary and ecological potential of yeast hybrids. Curr. Opin. Genet. Dev. 76, 101958 (2022).

15. Rinta-Harri, K. et al. Influence of Ploidy and Genetic Background on Stress Tolerance of Intraspecific Yeast Hybrids. Microb. Biotechnol. 19, e70337 (2026).

16. Dagilis, A. J., Kirkpatrick, M. & Bolnick, D. I. The evolution of hybrid fitness during speciation. PLoS Genet. 15, e1008125 (2019).

17. Bernardes, J. P., Stelkens, R. B. & Greig, D. Heterosis in hybrids within and between yeast species. J. Evol. Biol. 30, 538–548 (2017).

18. Schwarzkopf, E. J., Brandt, N. & Heil, C. S. The recombination landscape of introgression in yeast. eLife 13, (2024).

19. Morard, M. et al. Genomic instability in an interspecific hybrid of the genus *Saccharomyces*: a matter of adaptability. *Microb*. Genomics 6, e000448 (2020).

20. Bendixsen, D. P. et al. Reproductive Isolation due to Divergent Ecological Selection Is Accompanied by Vast Genomic Instability in Experimentally Evolved Yeast Populations. Mol. Ecol. 34, e70110 (2025).

21. Lancaster, S. M., Payen, C., Smukowski Heil, C. & Dunham, M. J. Fitness benefits of loss of heterozygosity in *Saccharomyces* hybrids. Genome Res. 29, 1685–1692 (2019).

22. Antunovics, Z., Nguyen, H.-V., Gaillardin, C. & Sipiczki, M. Gradual genome stabilisation by progressive reduction of the *Saccharomyces uvarum* genome in an interspecific hybrid with *Saccharomyces cerevisiae*. FEMS Yeast Res. 5, 1141–1150 (2005).

23. Toyomura, K. & Hisatomi, T. Postzygotic reproductive isolation among three *Saccharomyces* yeast species. Yeast 38, 326–335 (2021).

24. Greig, D., Borts, R. H., Louis, E. J. & Travisano, M. Epistasis and hybrid sterility in *Saccharomyces*. Proc. R. Soc. B Biol. Sci. 269, 1167–1171 (2002).

25. Zhang, Y. et al. Understanding the Nature of Hybrid Sterility and Divergence of Asian Cultivated Rice. Front. Plant Sci. 13, (2022).

26. Molecular Insights into Female Hybrid Sterility in Interspecific Crosses between *Drosophila melanogaster* and *Drosophila simulans*. https://www.mdpi.com/1422-0067/25/11/5681.

27. Ecological genomics of local adaptation | Nature Reviews Genetics. https://www.nature.com/articles/nrg3522.

28. Bundus, J. D., Wang, D. & Cutter, A. D. Genetic basis to hybrid inviability is more complex than hybrid male sterility in *Caenorhabditis nematodes*. Heredity 121, 169–182 (2018).

29. Fischer, G., James, S. A., Roberts, I. N., Oliver, S. G. & Louis, E. J. Chromosomal evolution in *Saccharomyces*. Nature 405, 451–454 (2000).

30. Brideau, N. J. et al. Two Dobzhansky-Muller Genes Interact to Cause Hybrid Lethality in *Drosophila*. Science 314, 1292–1295 (2006).

31. Fitzpatrick, B. M. Dobzhansky–Muller model of hybrid dysfunction supported by poor burst-speed performance in hybrid tiger salamanders. J. Evol. Biol. 21, 342–351 (2008).

32. Cutter, A. D. The polymorphic prelude to Bateson–Dobzhansky–Muller incompatibilities. Trends Ecol. Evol. 27, 209–218 (2012).

33. Lynch, M. & Force, A. G. The Origin of Interspecific Genomic Incompatibility via Gene Duplication. Am. Nat. 156, 590–605 (2000).

34. Alix, K., Gérard, P. R., Schwarzacher, T. & Heslop-Harrison, J. S. (Pat). Polyploidy and interspecific hybridization: partners for adaptation, speciation and evolution in plants. Ann. Bot. 120, 183–194 (2017).

35. Paun, O., Forest, F., Fay, M. F. & Chase, M. W. Hybrid speciation in angiosperms: parental divergence drives ploidy. New Phytol. 182, 507–518 (2009).

36. Kitano, J. & Okude, G. Causative genes of intrinsic hybrid incompatibility in animals and plants: what we have learned about speciation from the molecular perspective. Evol. J. Linn. Soc. 3, kzae022 (2024).

37. Lee, H.-Y. et al. Incompatibility of Nuclear and Mitochondrial Genomes Causes Hybrid Sterility between Two Yeast Species. Cell 135, 1065–1073 (2008).

38. Fishel, R. Mismatch Repair*. J. Biol. Chem. 290, 26395–26403 (2015).

39. Groothuizen, F. S. & Sixma, T. K. The conserved molecular machinery in DNA mismatch repair enzyme structures. DNA Repair 38, 14–23 (2016).

40. Pećina-Šlaus, N., Kafka, A., Salamon, I. & Bukovac, A. Mismatch Repair Pathway, Genome Stability and Cancer. Front. Mol. Biosci. 7, (2020).

41. Spies, M. & Fishel, R. Mismatch Repair during Homologous and Homeologous Recombination. Cold Spring Harb. Perspect. Biol. 7, a022657 (2015).

42. Elez, M. Mismatch Repair: From Preserving Genome Stability to Enabling Mutation Studies in Real-Time Single Cells. Cells 10, 1535 (2021).

43. Manhart, C. M. & Alani, E. Roles for mismatch repair family proteins in promoting meiotic crossing over. DNA Repair 38, 84–93 (2016).

44. Rogers, D. W., McConnell, E., Ono, J. & Greig, D. Spore-autonomous fluorescent protein expression identifies meiotic chromosome mis-segregation as the principal cause of hybrid sterility in yeast. PLoS Biol. 16, e2005066 (2018).

45. Goldfarb, T. & Alani, E. Distinct roles for the *Saccharomyces cerevisiae* mismatch repair proteins in heteroduplex rejection, mismatch repair and nonhomologous tail removal. Genetics 169, 563– 574 (2005).

46. Dujon, B. A. & Louis, E. J. Genome Diversity and Evolution in the Budding Yeasts (*Saccharomycotina*). Genetics 206, 717–750 (2017).

47. Rayssiguier, C., Thaler, D. S. & Radman, M. The barrier to recombination between *Escherichia coli* and *Salmonella typhimurium* is disrupted in mismatch-repair mutants. Nature 342, 396–401 (1989).

48. Worth, L., Clark, S., Radman, M. & Modrich, P. Mismatch repair proteins MutS and MutL inhibit RecA-catalyzed strand transfer between diverged DNAs. Proc. Natl. Acad. Sci. 91, 3238–3241 (1994).

49. Polosina, Y. Y., Mui, J., Pitsikas, P. & Cupples, C. G. The *Escherichia coli* Mismatch Repair Protein MutL Recruits the Vsr and MutH Endonucleases in Response to DNA Damage. J. Bacteriol. 191, 4041–4043 (2009).

50. Matic, I., Rayssiguier, C. & Radman, M. Interspecies gene exchange in bacteria: The role of SOS and mismatch repair systems in evolution of species. Cell 80, 507–515 (1995).

51. New, L., Liu, K. & Crouse, G. F. The yeast gene *MSH3* defines a new class of eukaryotic MutS homologues. Mol. Gen. Genet. MGG 239, 97–108 (1993).

52. Borts, R. H. et al. Mismatch repair-induced meiotic recombination requires the *pms1* gene product. Genetics 124, 573–584 (1990).

53. Datta, A., Adjiri, A., New, L., Crouse, G. F. & Jinks-Robertson, S. Mitotic Crossovers between Diverged Sequences Are Regulated by Mismatch Repair Proteins in *Saccharomyces cerevisiae*. Mol. Cell. Biol. 16, 1085–1093 (1996).

54. *MLH1*, *PMS1*, and *MSH2* Interactions During the Initiation of DNA Mismatch Repair in Yeast | Science. https://www.science.org/doi/10.1126/science.8066446.

55. Alani, E., Reenan, R. A. & Kolodner, R. D. Interaction between mismatch repair and genetic recombination in *Saccharomyces cerevisiae*. Genetics 137, 19–39 (1994).

56. Radman, M. & Wagner, R. Mismatch recognition in chromosomal interactions and speciation. Chromosoma 102, 369–373 (1993).

57. Bozdag, G. O. et al. Breaking a species barrier by enabling hybrid recombination. Curr. Biol. 31, R180–R181 (2021).

58. Greig, D. Reproductive isolation in *Saccharomyces*. Heredity 102, 39–44 (2009).

59. Ira, G., Malkova, A., Liberi, G., Foiani, M. & Haber, J. E. Srs2 and Sgs1–Top3 Suppress Crossovers during Double-Strand Break Repair in Yeast. Cell 115, 401–411 (2003).

60. De Muyt, A. et al. BLM helicase ortholog Sgs1 is a central regulator of meiotic recombination intermediate metabolism. Mol. Cell 46, 43–53 (2012).

61. Oh, S. D. et al. Sgs1, a Bloom’s helicase ortholog, prevents aberrant crossing-over by suppressing the formation of multi-chromatid joint molecules. Cell 130, 259–272 (2007).

62. Grandin, N. & Reed, S. I. Differential function and expression of *Saccharomyces cerevisiae* B-type cyclins in mitosis and meiosis. Mol. Cell. Biol. 13, 2113–2125 (1993).

63. Cross, F. R., Yuste-Rojas, M., Gray, S. & Jacobson, M. D. Specialization and Targeting of B-Type Cyclins. Mol. Cell 4, 11–19 (1999).

64. Guan, Y., Dunham, M. J., Troyanskaya, O. G. & Caudy, A. A. Comparative gene expression between two yeast species. BMC Genomics 14, 33 (2013).

65. Loidl, J., Jin, Q.-W. & Jantsch, M. Meiotic pairing and segregation of translocation quadrivalents in yeast. Chromosoma 107, 247–254 (1998).

66. Forejt, J. & Jansa, P. Meiotic Recognition of Evolutionarily Diverged Homologs: Chromosomal Hybrid Sterility Revisited. Mol. Biol. Evol. 40, msad083 (2023).

67. Colson, I., Delneri, D. & Oliver, S. G. Effects of reciprocal chromosomal translocations on the fitness of *Saccharomyces cerevisiae*. EMBO Rep. 5, 392–398 (2004).

68. Hou, J., Friedrich, A., de Montigny, J. & Schacherer, J. Chromosomal Rearrangements as a Major Mechanism in the Onset of Reproductive Isolation in *Saccharomyces cerevisiae*. Curr. Biol. 24, 1153–1159 (2014).

69. Ono, J. & Greig, D. A *Saccharomyces* paradox: chromosomes from different species are incompatible because of anti-recombination, not because of differences in number or arrangement. Curr. Genet. 66, 469–474 (2020).

70. Hung, Y.-L. et al. Rad51, Rad54, and ZMM proteins antagonize the mismatch repair system to promote fertility of budding yeast intraspecies hybrid zygotes. Nucleic Acids Res. 53, gkaf847 (2025).

71. Mozzachiodi, S. et al. Aborting meiosis allows recombination in sterile diploid yeast hybrids. Nat. Commun. 12, 6564 (2021).

72. Naseeb, S. et al. Restoring fertility in yeast hybrids: Breeding and quantitative genetics of beneficial traits. Proc. Natl. Acad. Sci. U. S. A. 118, e2101242118 (2021).

73. McNeill, J., Brandt, N., Schwarzkopf, E. J., Jimenez, M. & Smukowski Heil, C. Impacts of temperature on recombination rate and meiotic success in thermotolerant and cold-tolerant yeast species. Heredity 134, 473–484 (2025).

74. Martínez, A. A. & Lang, G. I. Pervasive weak incompatibilities shape phenotypic variation and genome evolution in yeast hybrids. Nat. Commun. 17, 10662 (2026).

75. Gietz, R. D. & Schiestl, R. H. High-efficiency yeast transformation using the LiAc/SS carrier DNA/PEG method. Nat. Protoc. 2, 31–34 (2007).

76. Samsonova, I. A., Böttcher, F., Steinwehr, D. & Schilowa, B. A simple technique for random spore analysis of yeasts using nystatin. J. Microbiol. Methods 3, 283–290 (1985).

77. Tong, A. H. et al. Systematic genetic analysis with ordered arrays of yeast deletion mutants. Science. 294. 2364–8 (2001).

78. Dunham, M. J., Gartenberg, M. R., & Brown, G. W. Methods in yeast genetics and genomics: a Cold Spring Harbor Laboratory course manual. Cold Spring Harbor Laboratory Press (2015).

