## Supplemental file 1 for "Meiotic null *MSH2* and *SGS1* alleles in *S. cerevisiae* x *S. uvarum* hybrids result in near-haploid offspring with mixed parental chromosomal inheritance"

### Slide 1
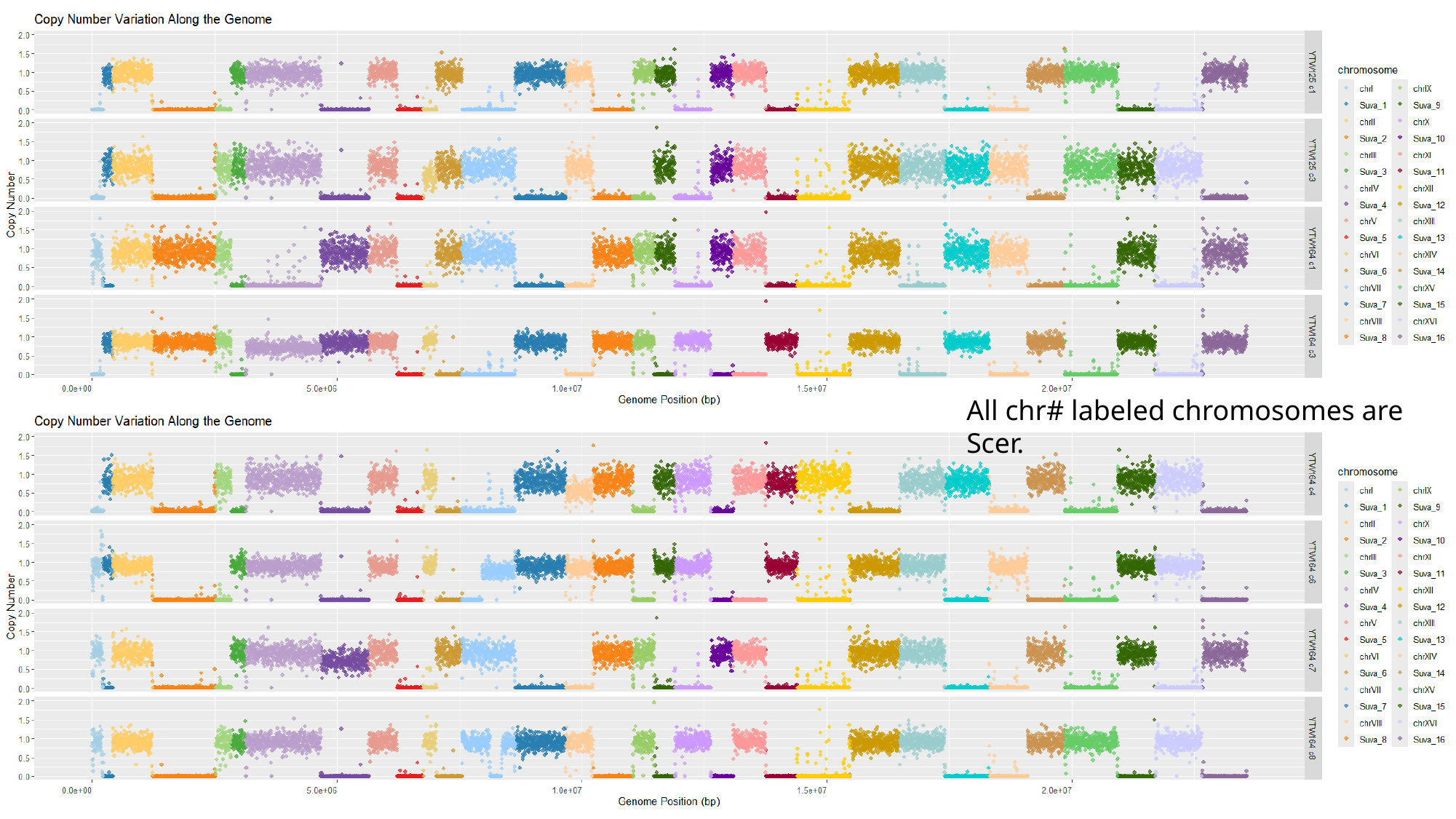

All chr# labeled chromosomes are Scer.

### Slide 2
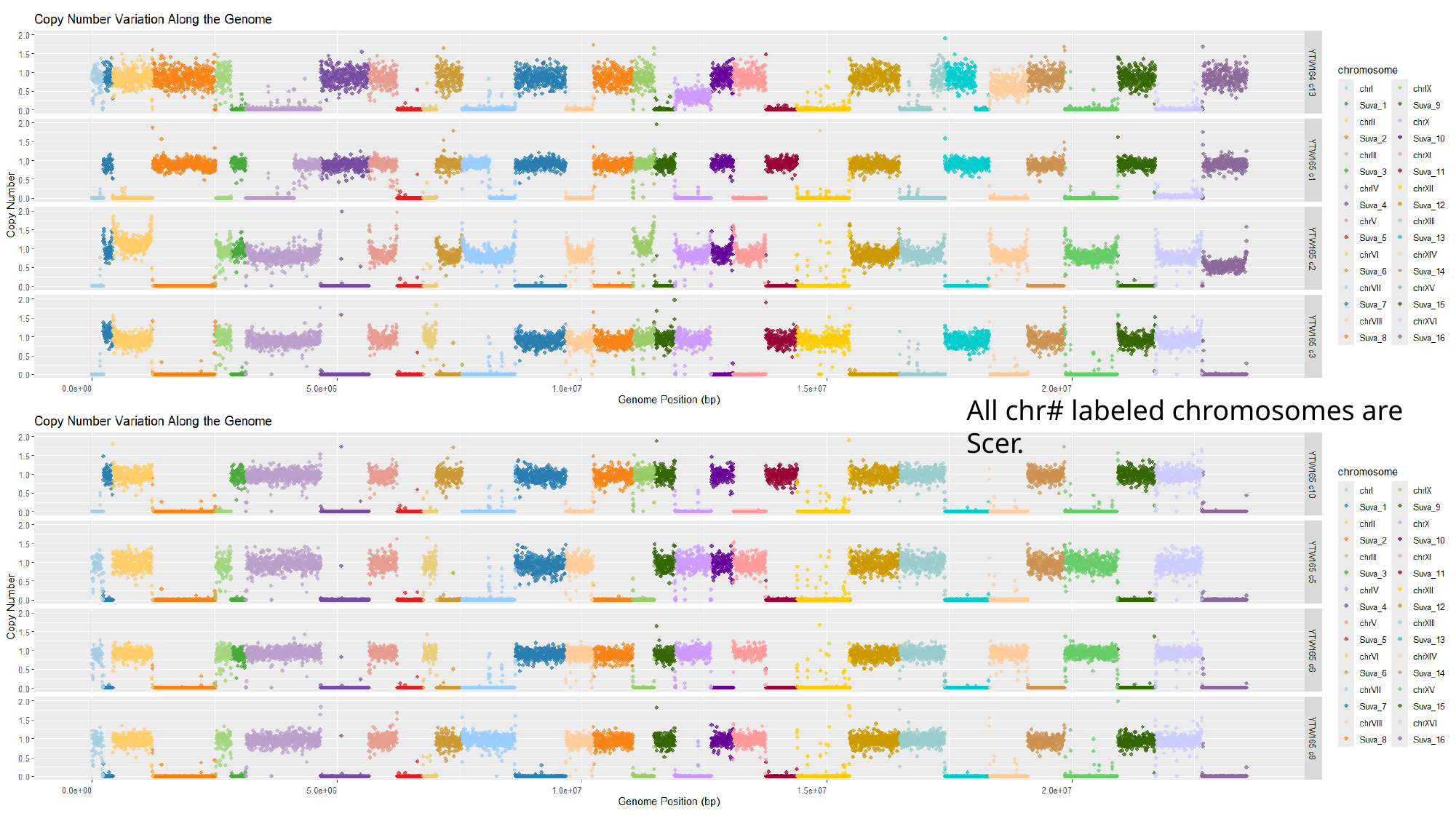

All chr# labeled chromosomes are Scer.

### Slide 3
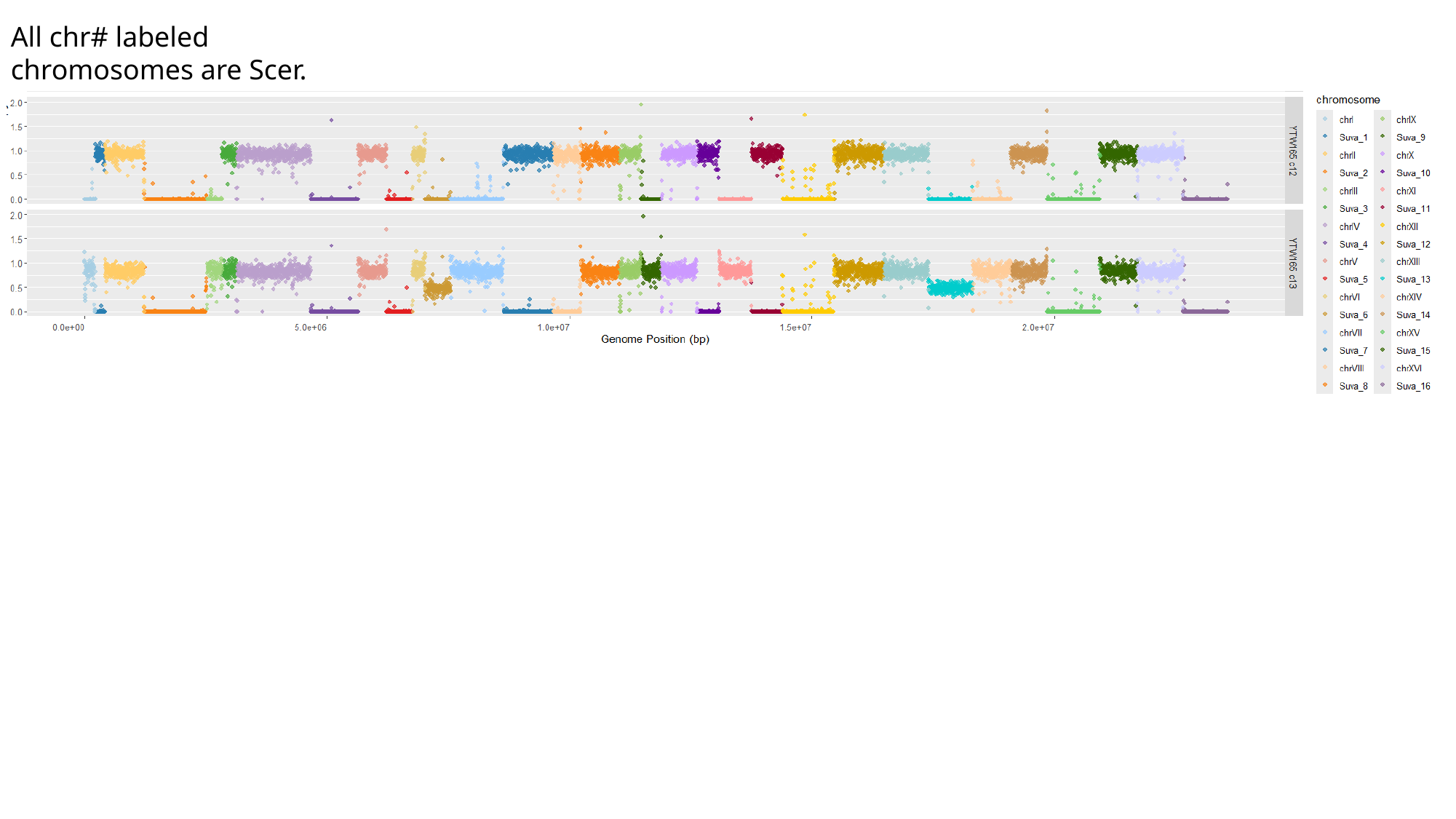

All chr# labeled chromosomes are Scer.
